# Sessile bacterium unlocks ability of surface motility through mutualistic interspecies interaction

**DOI:** 10.1101/2020.04.24.060913

**Authors:** Miaoxiao Wang, Shuang Geng, Bing Hu, Yong Nie, Xiao-Lei Wu

## Abstract

In addition to their common planktonic lifestyle, bacteria frequently live in surface-associated habitats. Surface motility is essential for exploring these habitats for food sources. However, many bacteria are found on surfaces, even though they lack features required for migrating along surfaces. How these canonical non-motile bacteria adapt to the environmental fluctuations on surfaces remains unknown. Recently, several cases of interspecies interaction were reported that induce surface motility of non-motile bacteria either by using ‘hitchhiking’ strategies or through ‘social spreading’ mechanisms. Here, we report a previously unknown mechanism for interaction-dependent surface motility of the canonical non-motile bacterium, *Dietzia* sp. DQ12-45-1b, which is induced by interaction with a dimorphic prosthecate bacterium, *Glycocaulis alkaliphilus* 6B-8^T^. *Dietzia* cells exhibits “sliding”-like motility in an area where the strain *Glycocaulis* cells was pre-colonized with a sufficient density. Furthermore, we show that biosurfactants play a critical role in inducing the surface motility of *Dietzia* cells. Our analysis also demonstrates that *Dietzia* degrade n-alkanes and provide *Glycocaulis* with the resulting metabolites for survival, which in turn enabled directional migration of *Dietzia* towards nutrients in the environment. Such interaction-dependent migration was also found between *Dietzia* and *Glycocaulis* strains isolated from other habitats, suggesting that this mutualistic relationship ubiquitously occurs in natural environments. In conclusion, we propose a novel model for such a ‘win-win’ strategy, whereby non-motile bacteria pay metabolites to dimorphic prosthecate bacteria in return for migrating to reach environments otherwise inaccessible. We propose that this mechanism represents a common strategy for canonically non-motile bacteria living on a surface.

**Importance:** Cell motility provides a selective advantage for bacteria searching for nutrients. While a large body of evidence exists for how motile bacteria migrate on surface by virtue of different ways of motility, fewer studies concerned about how canonical non-motile bacteria adapted to those surface-associated habitats. Recent reports have proposed that interactions with other bacteria trigger the movement of those sessile bacteria. However, these interactions are limited to ‘hitchhiking’ or ‘social spreading’ modes. Here, we characterized a previously unknown interaction mode between *Dietzia* and *Glycocaulis*.

This interaction differs from previously described modes, thus advance our limited understanding of how sessile bacteria move on surfaces. We propose that this interaction mode represents a ‘win-win’ strategy for both strains, and this mode might be widely distributed across diverse environments. These novel insights should greatly assist in understanding the mechanisms responsible for the cellular interplay between microbes in complex microbiomes.

## Introduction

In addition to growing in a planktonic form, many bacteria also colonize surfaces, a process that requires attachment of bacterial cells to various surfaces and surface-associated motility (1–3). A large variety of such surface motility behaviors have been identified, including swarming mediated by flagella, twitching and gliding mediated by pili, as well as passive sliding created by expansive forces derived from colony growth (3). These movements allow bacteria to escape local stresses and explore their environment in search for nutrients (1, 3, 4). Motility is also involved in other bacterial processes occurring on surfaces, including biofilm formation, microbe-host interaction, and bacterial cell morphogenesis (1, 3, 4). However, while these motility-related traits provide considerable selective advantage for bacteria living on surface, many bacterial species, called canonically non-motile bacteria, lack such migratory characteristics. How these bacteria adapt to heterogeneous environments remains to be elucidated (5, 6).

The social life in complex communities offers opportunities for bacterial cells to interact with other populations. Benefiting from such interactions, a number of canonically non-motile bacteria have been described to move on surface by interacted with other social groups. For example, ‘riding’ on the swarming *Paenibacillus vortex* cells promote the dispersal of non-motile *Xanthomonas* (7) as well as *Escherichia coli* (8) strains on solid surfaces. Gliding motility of *Capnocytophaga gingivalis* population plays an import role in long-range transport of those non-motile bacterial cells in a polymicrobial community (9). This type of social behavior also contributes to shaping of spatial patterns of multi-species bacterial colonies (10). Together, these phenomena are called ‘hitchhiking’ strategies (6), which requires a highly motile population acting as a ‘truck’ carries non-motile ‘cargo’ cells along. In addition, a recent case showed that two sessile soil strains, *Pseudomonas fluorescens* Pf0-1 and *Pedobacter* sp. V48 started co-migrating upon initial close association (11). However, due to the diversity of social interactions in microbial world, it still remains an open question whether there are other interaction modes that can trigger the surface motility of canonically non-motile bacteria.

Here we report a previously unknown form of surface motility way of non-motile bacteria, *Dietzia* spp. that was induced by dimorphic prosthecate bacteria (DPB), *Glycocaulis* spp., via interspecies contact-dependent interactions. The underlying mechanism of this interaction is different from the previous reported manners. The presence of the DPB induced the *Dietzia* species migrate outward on a soft agar surface and form a dendritic fractal pattern, while the DPB earned essential carbon sources from the partner and survive better in the nutrient-limited environment. Therefore, this interaction mode is a ‘win-win’ strategy for both strains to live on surface-attached habitats.

## Results

### Interspecies interaction induces bacterial surface motility

*Dietzia* sp. DQ12-45-1b (45-1b) and *G. alkaliphilus* 6B-8^T^ (6B-8) were both isolated from one oil reservoir in the Daqing Oilfield, China [9, 10]. Strain 45-1b is a Gram-positive, canonically non-motile bacterium with the ability to biodegrade a wide range of petroleum hydrocarbons, especially showing a great ability of utilizing *n*-alkanes with a wide range of chain length (12). Strain 6B-8 is a Gram-negative bacterium that can grow in Luria Broth (LB) medium and does not show typically surface motility (Fig. 1a) on LB plate with 0.5% agar. Our previous analysis also suggests that 6B-8 only utilizes a very narrow spectrum of carbon sources, excluding both alkanes and glucose (13).

**Figure 1.**
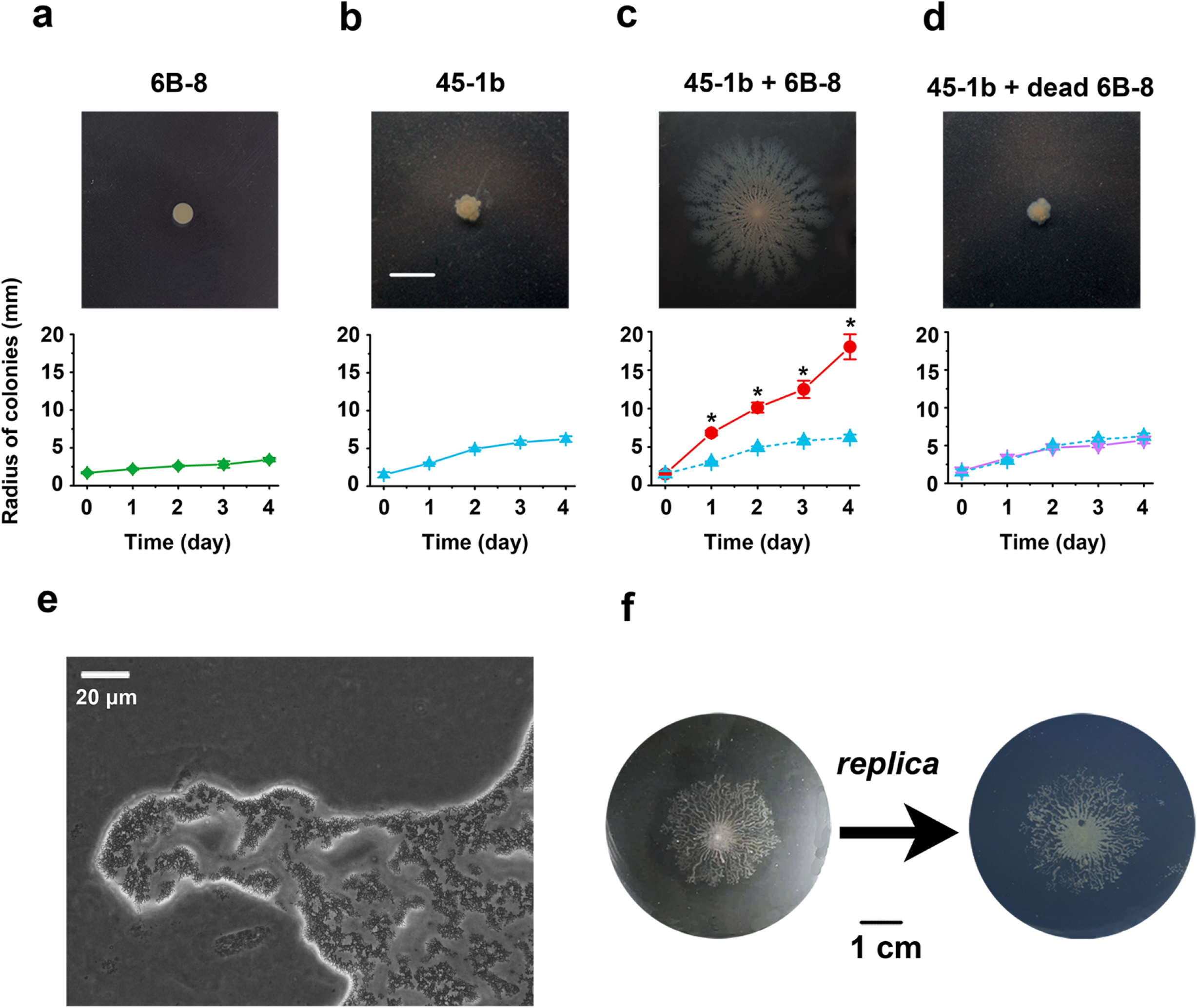
Surface motility of *Dietzia* sp. DQ12-45-1b induced by *Glycocaulis alkaliphilus* 6B-8^T^. Movement of 6B-8 (**a**) on a LB plate containing 0.5% agar and 45-1b on alkane agar plate in the presence of live (**c**), dead (**d**), or absence of (**b**) 6B-8. The lower panel depicts temporal changes in colony radii. Data represent mean ± SD of three independent replicates. Dashed lines in (**c**) and (**d**) indicate changes in colony radii in (**a**) and statistical analyses were done by Student’s t-test: *, *p* < 0.01. Colony morphology **(e)** shows that 45-1b cells form cell aggregates at the edge of the colony and move together during migration. Microscopic images were obtained from the edge of the dendritic fractal colony. **(f)** Replica plate analysis shows that 6B-8 cells grow at sites where 45-1b cells grow on alkane agar plates. The left image shows the colony morphology of 45-1b on soft agar pre-mixed with 6B-8 using C_16_ as the sole carbon source. The right image shows the colony morphology of 6B-8^T^ cells after blotting on LB agar containing 5 μg/mL kanamycin.

As expected, while the strain 45-1b formed a smooth circular and static colony on 0.5% agar plates (Fig. 1b) containing hexadecane (C_16_) as the sole carbon source (henceforth called ‘alkane agar plate’), the strain 6B-8 failed to grow alone under these conditions. Intriguingly, when the strain 45-1b suspension was dropped on the same agar plate (containing C_16_ as the carbon source) thoroughly premixed with 6B-8 cells, we found that the 45-1b cells developed a fractal dendritic colony with a colony spreading speed of 2.87 ± 0.28 μm/min (Fig. 1c). Together, these results suggest that 6B-8 cells assist the 45-1b cells in migrating along the surface, thus exhibiting a behavior characteristic of ‘surface motility’. This surface motility was observed only when live 6B-8 cells were present (Fig. 1d), suggesting that the movement of the strain 45-1b critically depended on the interaction with 6B-8. Analysis by light microscopy showed that the cells of 45-1b formed cell aggregates at the edge of the colony and migrated in a synchronous manner (Fig. 1e).

To investigate the location of 6B-8 interacting with 45-1b in the agar plate with C_16_, the entire plate was blotted using a 0.22-μm sterile membrane and transferred to a new LB agar plate containing kanamycin. As only the strain 6B-8 is kanamycin-tolerant, the colonies we detected on the kanamycin-containing plate were identified as belonging to strain 6B-8. Interestingly, the colonies of 6B-8 that formed on the agar were identical to the fractal dendritic spreading path of the 45-1b colonies on the agar plate (Fig. 1f), which demonstrated that the pre-mixed 6B-8 cells grew at the sites at which 45-1b grew in the agar plate with C_16_ as the carbon source. Together, these results indicated that the cells of strain 45-1b were able to move on the surface via interacting with cells of strain 6B-8.

### *Dietzia* sp. DQ12-45-1b only migrated to the region pre-colonized by *G. alkaliphilus* 6B-8^T^

A number of recent studies reported that highly motile bacterial populations can transport sessile populations as cargo on surface (7–10). We therefore investigated whether strain 45-1b was also transported as the cargo by 6B-8. To this end, we performed a surface motility assay similar to that described in a previous study (8). In brief, we added an inoculum containing an equal mixture of 6B-8 and 45-1b to the agar plate and co-cultivated them with C_16_ as the sole carbon source. Unexpectedly, we found that 45-1b failed to move on the agar surface (Fig. 2a), indicating that the surface motility of 45-1b was not induced by the movement of 6B-8. Instead, when we spread 6B-8 cells out on the agar plates to form expected patterns (e.g., semi-circle or cross), and inoculated 45-1b suspension on the center of these patterns, we found that the strain 45-1b developed colonies only within the regions where 6B-8 was previously overlaid (Fig. 2b). Together, these results indicated that the 45-1b cells only migrated to where 6B-8 cells were present.

**Figure 2.**
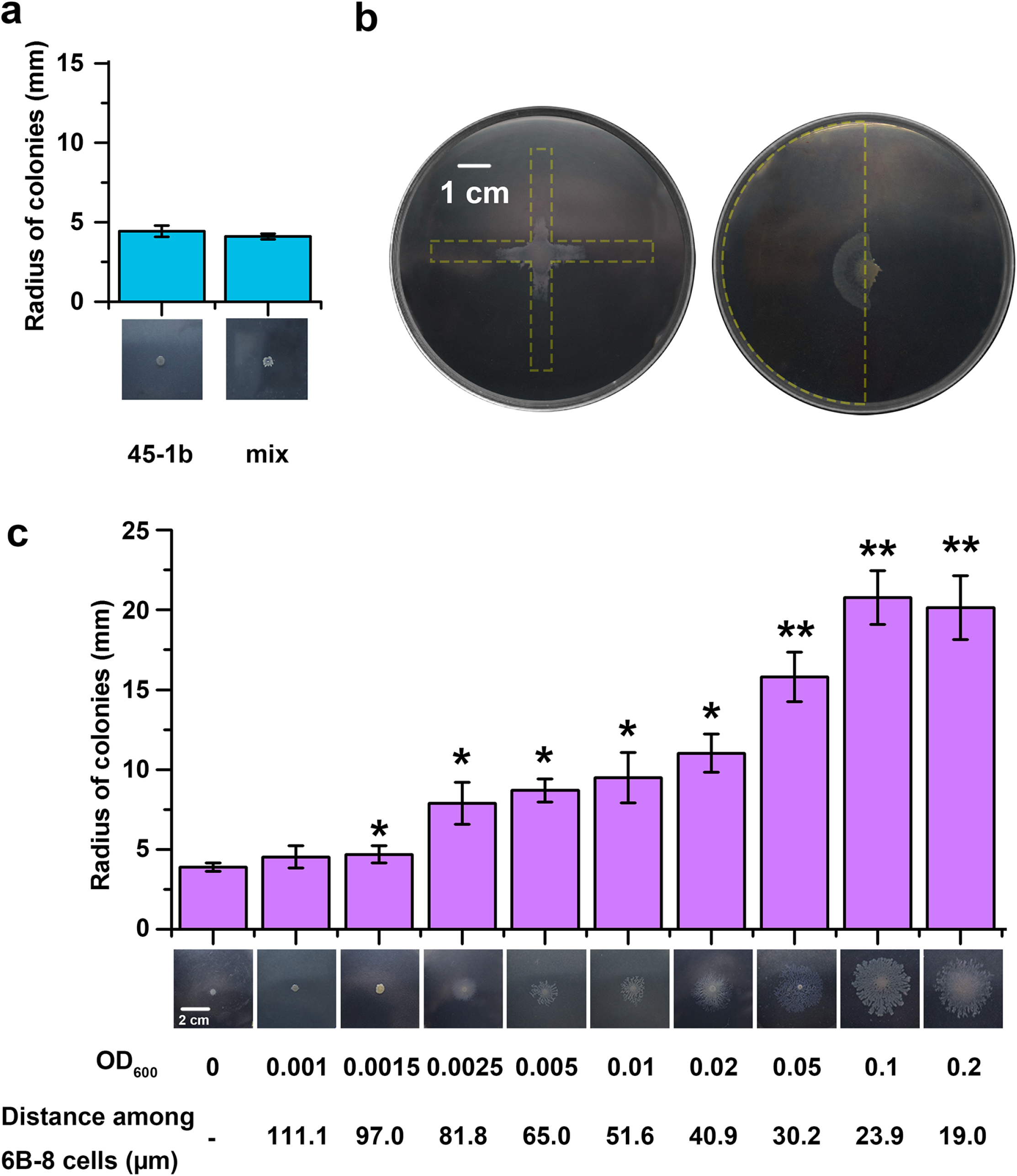
*Dietzia* sp. DQ12-45-1b only migrated to the region pre-colonized by *G. alkaliphilus* 6B-8^T^. **(a)** When the two strains were initially mixed together at equal amounts and inoculated onto a soft agar plate, no significant surface motility was observed; Statistical analysis by t-test compared with 45-1b (the movement of 45-1b in the absence of 6B-8), *p* = 0.27. **(b)** Colonies of 45-1b cells only ‘moved’ at locations where 6B-8 cells were previously overlaid. The left image shows 6B-8 cells previously overlaid within the cross region indicated by dashed lines. The right image shows 6B-8 cells previously overlaid within the left half of the agar, as indicated by dashed lines. **(c)** Strain 45-1b cells exhibit surface motility only when a set number of 6B-8 cells was initially colonized the outward region. The OD_600_ values reflect the initial concentrations of 6B-8 cells premixed in the soft agar. The distance values reflect the estimated distances between 6B-8 cells when they were premixed in soft agar, which were calculated using a cubic lattice model (see Methods for detail). These experiments were carried out providing C_16_ as the sole carbon source. Data shown in (**a)** and (**c)** represent three independent experiments; the error bars represent standard deviation. Statistical analysis was done by Student’s t-test: *, *p* < 0.05 and **, *p* < 0.01.

To further investigate the effect of initial spatial distribution of 6B-8 on the surface motility of 45-1b, we incubated cells of strain 45-1b in agar plates premixed with different starting 6B-8 cell densities, using C_16_ as the sole carbon source. To achieve different degrees of spatial separations, we used 6B-8 cell densities ranging from OD_600_ values of 0.001 to 0.2, as a result of which the average distances among 6B-8 cells ranged from 111 μm to 19 μm in the agar plates (as calculated using a cubic lattice model, see Methods for more details). The results showed that when the mean distance among pre-mixed 6B-8 cells was shorter than 97 μm, 45-1b colonies exhibited a clearly visible dendritic fractal pattern. As shown in Fig. 2c, the apparent colony spreading speed was between 1.37 ±0.23 and 3.50 ±0.35 μm/min. Taken together, these results showed that the migration of the strain 45-1b required a set number of 6B-8 cells initially spatially separated from 45-1b, and the mean distance between each pair of 6B-8 cells had to be shorter than 97 μm. These observations differ completely from the previously reported ‘hitchhiking’ strategy between non-motile and highly motile bacteria (7–10). In addition, the interaction modes of 45-1b and 6B-8 are also distinct compared to those of the two sessile strains, *Pseudomonas fluorescens* Pf0-1 and *Pedobacter* sp. V48 (11), suggesting different underlying mechanisms for interaction-dependent surface motility.

### Surface motility of *Dietzia* sp. DQ12-45-1b required direct contact with *G. alkaliphilus* 6B-8^T^

We next asked whether cells of the strain 6B-8 trigger the movement of 45-1b cells by way of metabolite exchange between the two cell types. Firstly, we used a Transwell plate in which the two strains were separated by a microporous barrier membrane that allowed exchange of their extracellular metabolites, verified by a fluorescein diffusion assay (Fig. 3a). After incubation with C_16_ as the sole carbon source, no surface motility was observed (Fig. 3a). Furthermore, we failed to observe any surface motility behavior when the two strains were separated on an agar slide by either 1 or 2 cm (Fig. 3b, left), a distance at which metabolites can also diffuse well as fluorescein (Fig. 3b, right). Next, we inoculated cells of both 45-1b and 6B-8 strains into 150 mL minimal medium containing C_16_ as the sole carbon source, and incubated at 30°C. Subsequently, we removed all cells by centrifuging and filtering, and harvested the supernatant containing metabolites secreted by both 45-1b and 6B-8. We then incubated the cells of strain 45-1b on an agar plate that was overlaid with the co-culture supernatant. As shown in Fig. 3c, no surface motility was observed. Together, these results revealed that metabolite exchange was not sufficient to induce surface motility in strain 45-1b.

**Figure 3.**
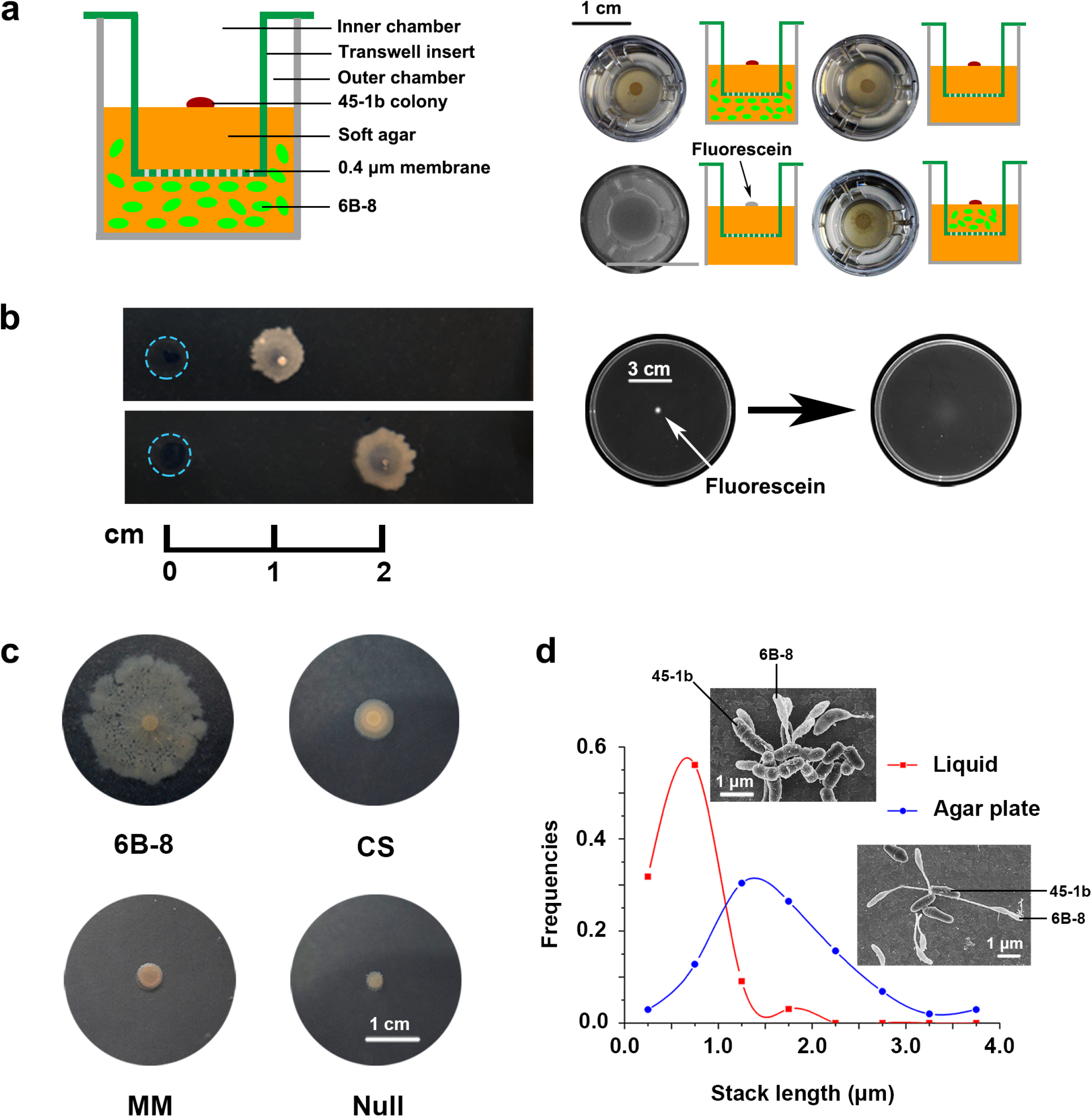
Surface motility of *Dietzia* sp. DQ12-45-1b required direct contact with *alkaliphilus* 6B-8^T^. (**a**) Incubation of the two strains separated by a 0.4-μm transwell membrane barrier, allowing metabolite exchange between the inner and outer wells, without cell migration across the barrier. (**b**) Incubating the two strains at distances of 1 cm and 2 cm on a 0.5% agar plate failed to induce surface motility of 45-1b. (**c**) Incubation of 45-1b with co-incubation supernatant of both strains failed to induce motility. The soft agar plates were overlaid with the co-incubation supernatant or minimal medium and then used for surface motility assay. CS: co-incubation supernatant; MM: minimum medium; Null: without any addition. C_16_ was used as the sole carbon source in these three cases. (**d**) 6B-8 cells stretched their stalks differently to attach to 45-1b cells when they were co-cultured on soft agar (upper) and in liquid medium (lower) using hexadecane as the sole carbon source. 6B-8 cells had significantly longer (t-test, *p* < 0.01) stalk lengths when cultured on soft agar than in liquid medium.

To investigate the potential interaction via direct cell-cell contact between the two strains, we collected samples from the migration areas of 45-1b, and performed scanning electron microscopy (SEM) observation. The images showed that 6B-8 cells attached to the 45-1b cells using their stalks, suggesting that the cells of these two strains interacted via cell-cell contacts. Moreover, the length of the stalks in this scenario was significantly longer than the stalks of 6B-8 cells harvested from co-culture of the two strains in liquid medium with shaking (Fig. 3d). Together, these results suggested that a direct contact interaction contributes to the induction of surface motility in strain 45-1b cells.

### Surface motility of *Dietzia* sp. DQ12-45-1b requires surfactants produced from alkane metabolism

To test whether the surface motility in 45-1b co-incubated with 6B occurs under various environmental conditions, we incubated 45-1b cells on agar plates premixed with 6B-8 cells, each plate containing one of 23 different compounds as sole carbon source (Table 2). While 45-1b cells can metabolize all of the 23 compounds, 6B-8 cells only metabolize acetic acid and α-ketoglutarate. Surprisingly, upon co-incubation with 45-1b cells on agar plates, we observed growth of 6B-8 cells for all 23 carbon sources, indicating that the growth of 6B-8 cells was supported by the metabolic by-products of 45-1b cells. However, we only found surface motility of 45-1b cells in agar plates containing three alkanes (i.e. dodecane, tetradecane and hexadecane; Table 2; Fig. 4a). This indicates that the surface motility of 45-1b in this interaction depends on specific carbon-sources.

**Figure 4.**
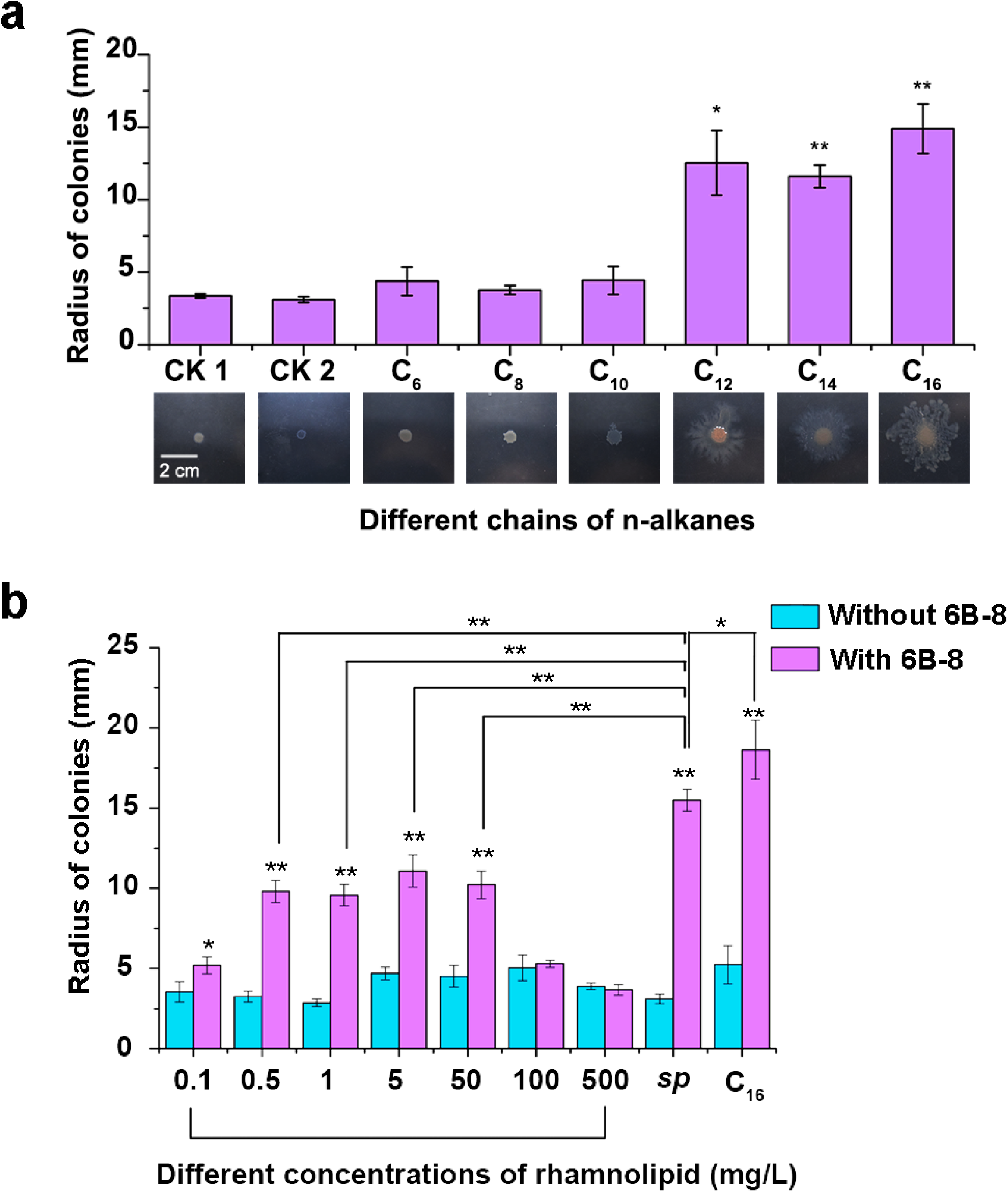
Surface motility of *Dietzia* sp. DQ12-45-1b required surfactants produced from alkane metabolism. **(a)** Effect of the carbon chain length of *n*-alkanes on induction of 45-1b surface motility. Data represent mean± SD of three independent replicates. The lower panel shows the colony pattern of 45-1b when co-cultured with 6B-8 using the corresponding carbon sources. CK1, in the absence of 6B-8 and using C_16_ as the carbon source. CK2, in the presence of 6B-8 but without carbon source addition. Statistical analysis was performed by comparison with CK1 using Student’s t-test: *, *p* < 0.05, and **, *p* < 0.01. **(b)** Induction of 45-1b surface motility by bio-surfactant concentrations ranging from 0.1 to 50 mg/L and *sp*: surfactant-containing supernatant. For incubation with rhamnolipid or *sp*, a 45-1b cell suspension was inoculated onto soft agar pre-mixed with 6B-8 cells and glucose as the sole carbon source. For C_16_ (positive control), a 45-1b cell suspension was inoculated onto soft agar pre-mixed with 6B-8 cells with C_16_ as the sole carbon source.

In one of our previous studies, we reported that upon growth on n-alkanes, 45-1b cells produce biosurfactants to emulsify alkanes and achieve surfactant-mediated access (14). Moreover, the presence of biosurfactants has been reported to facilitate bacterial surface motility (1, 5). Therefore, we hypothesized that the surfactants produced from alkane metabolism are necessary for the surface motility of 45-1b cells. To test this hypothesis, we first acquired the biosurfactant-containing supernatant by mono-culturing of 45-1b cells in liquid minimal medium containing C_16_ as the sole carbon source. We then incubated 45-1b on the agar plate both premixed with 6B-8 and overlaid the biosurfactant-containing supernatant on the surface, using glucose as the sole carbon source. Since glucose was not sufficient to trigger the interaction-dependent surface motility of 45-1b (Table 2), we wondered whether a turnover would occur in the presence of the biosurfactant-containing supernatant. As shown in Fig. 4b (denoted by ‘*sp*’), surface motility of 45-1b was observed. To confirm that this turnover was due to the effects of the secreted biosurfactant, we overlaid solutions of rhamnolipid on the surface, and performed the similar assays. As shown in Fig. 4b, surface motility of 45-1b cells was observed with concentrations of rhamnolipid ranging from 0.1 to 50 mg/L (Fig. 4b). Together, these results indicated that surface motility of *Dietzia* 45-1b requires biosurfactants produced from its alkane metabolism.

### Both cell types yield from the interaction

To analyze the benefits 6B-8 cells gain from this interaction with 45-1b cells, we compared the biomass of 6B-8 either co-cultured with 45-1b or monocultured on agar surface by the replica blotting assay. In the co-culturing assays, after blotting, the 6B-8 colony showed a similar dendritic fractal pattern that exactly matched the colonies of 45-1b on the alkane agar plate (Fig. 1f). In contrast, when 6B-8 was premixed with agar plates without incubation with 45-1b cells, we found that much less visible colonies had formed after blotting (Fig. 5a). Together, these results indicated that the strain 45-1b utilized C_16_ and simultaneously provided 6B-8 cells with essential nutrients, presumably for survival and/or growth. To further confirm this idea, we overlaid supernatants from a monoculture of 45-1b cells in C_16_ liquid medium or α-ketoglutarate, a derivative produced during C_16_ biodegradation by 45-1b, on the surface of agar plate premixed with 6B-8 but without inoculating 45-1b. After same replica blotting operation, more colonies of grew on the LB agar plate containing kanamycin than scenarios without addition of metabolites (Fig. 5a). Together, these results indicated that the 6B-8 cells obtained the nutrients essential for survival and/or growth through metabolic interaction with 45-1b cells.

**Figure 5.**
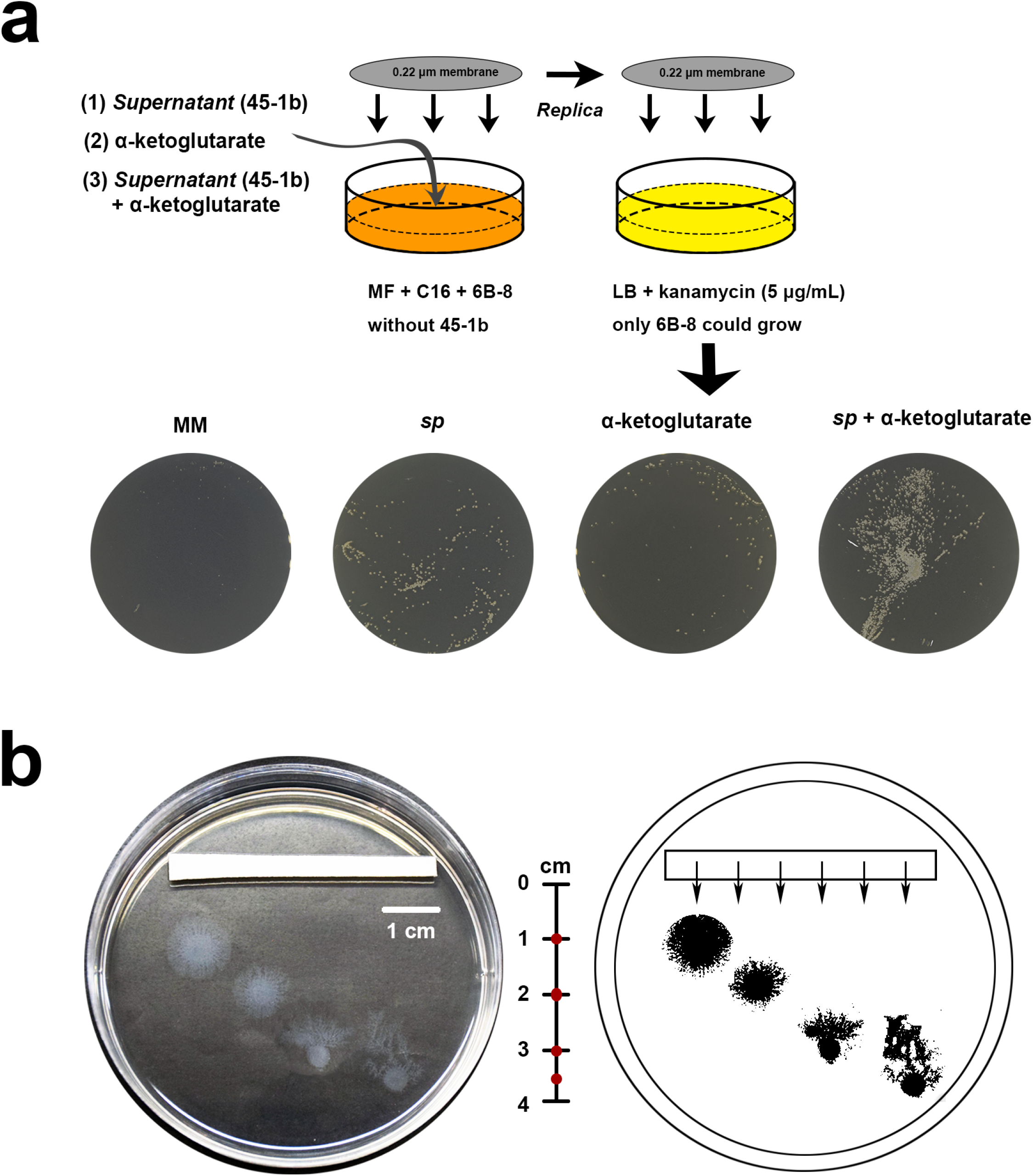
Strain interaction is mutually beneficial. **(a)** 6B-8 cells survived better on an agar plate overlaid with the metabolite-containing supernatant (*sp*) or α-ketoglutarate, a metabolite of 45-1b degraded hexadecane. The agar plate overlaid with the minimum medium (MM) was used as control. Three independent replicated experiments were conducted, representative results are shown in the lower panel, depicting the colonies of 6B-8 growing on the LB agar plates. These results suggested that the metabolites produced by 45-1b assist 6B-8 cell survival /or growth. **(b)** When grown in a nutrient gradient environment containing 6B-8 cells, 45-1b grew and migrated towards high concentrations of nutrients. Soft agar plates were pre-mixed with 6B-8 cells; biosurfactant-containing supernatant was spread on this plate to induce surface motility, a strip of sterile paper (4 cm×0.35 cm) saturated with 20 μL of a glucose solution (250 g/L) was placed on one side of the plate to form a nutrient gradient. Strain 45-1b was inoculated by dropping it on the surface of the soft agar. The ruler indicates the distance from the paper. Red dots represent the positions where 45-1b was inoculated. The binary image on the right shows the direction of surface motility. The arrows indicate the direction of the nutrient gradient.

In many bacterial species, cell motility is associated with the search for nutrients. To better understand whether increased motility of 45-1b was associated with seeking nutrients, we developed a glucose gradient on an agar plate that was pre-mixed with 6B-8 cells and overlaid with the biosurfactant-containing supernatant, followed by inoculation of *Dietzia* 45-1b. As shown in Fig. 5b, 45-1b displayed chemotaxis towards the higher glucose concentration. This result suggested that, by interacting with 6B-8 cells, the 45-1b cells were able to move towards environments containing more nutrient (i.e. chemotaxis). Taken together, our results indicate that both cell types yield from this interaction, resulting in a ‘win-win’ situation on surface-attached habitats.

### The interaction mode is widespread across many ecological habitats

In order to assess whether this interaction was specific for these two strains, or whether this is a more general phenomenon, we first tested whether other bacteria or related dimorphic prosthecate bacteria can assist in the migration of 45-1b cells. In these assays, *Glycocaulis abyssi* MCS33 (isolated from a deep-sea hydrothermal vent) and *Glycocaulis albus* SLG210-30A1^T^ (isolated from oil-contaminated saline soil) also induced surface motility for 45-1b cells (Fig. 6a). Moreover, 6B-8 cells also induced surface motility of *Dietzia psychralcaliphila* ILA-1^T^ (isolated from a drain of a fish product processing plant) and *Dietzia timorensis* DSM 45568^T^ (isolated from soil; Fig. 6b). As these strains were isolated from a variety of environments, we propose that this interaction mode is present across a wide range of ecological habitats.

**Figure 6.**
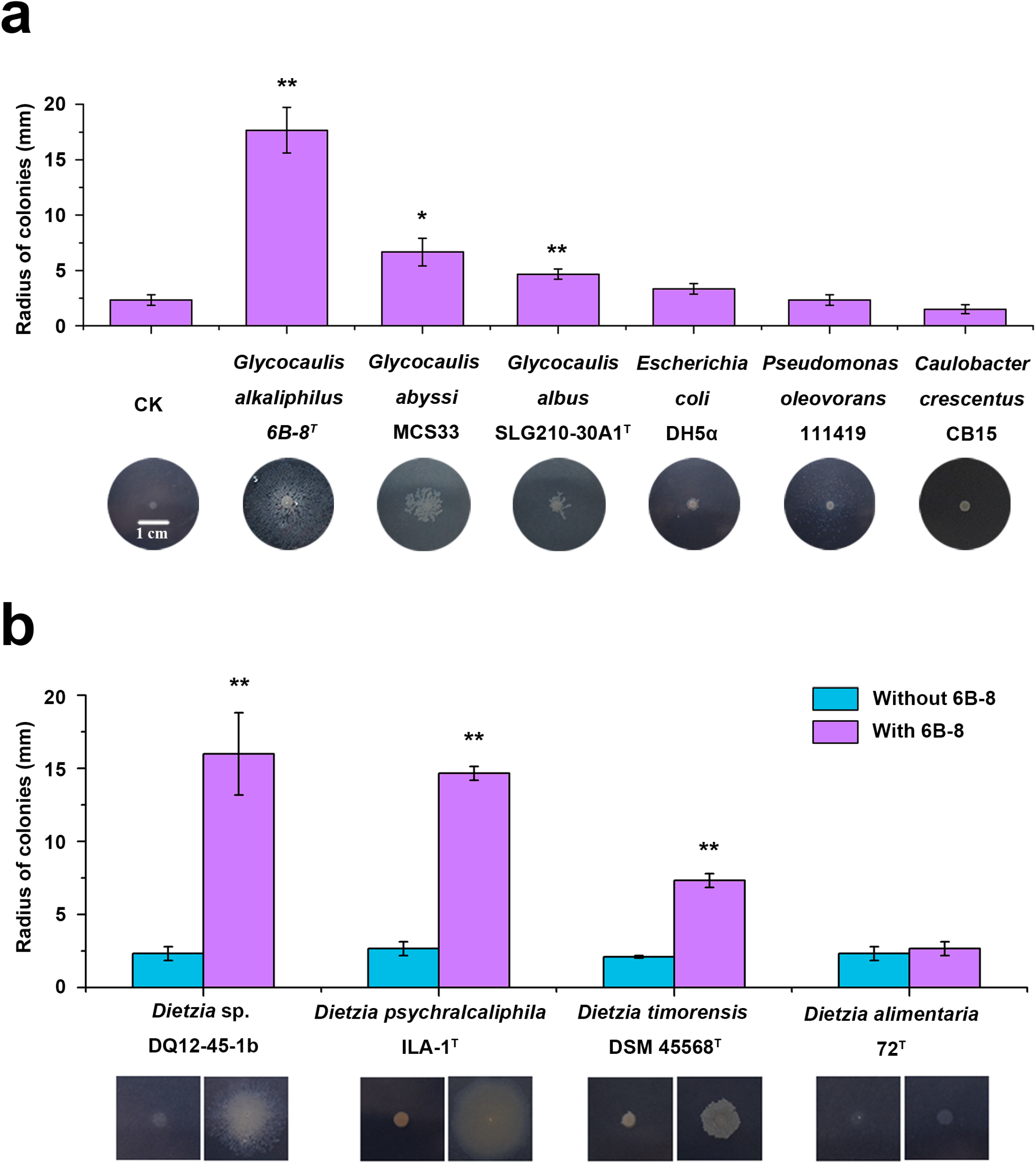
Surface motility is induced in different *Dietzia* strains, and in response to various *Glycocaulis* strains. (**a**) Only *Glycocaulis* strains were able to induce *Dietzia* surface motility. The DPB strain, *Caulobacter* sp. CB15 failed to induce *Dietzia* surface motility, indicating that *Glycocaulis* and *Caulobacter* use different survival strategies to adapt to environments. CK, the movement of *Dietzia* 45-1b in the absence of *Glycocaulis* 6B-8^T^. (**b**) Other *Dietzia* strains also exhibited induction of surface motility by *G. alkaliphilus*, except for *Dietzia alimentaria* 72^T^, suggesting that induced surface motility may be universal to *Dietzia* strains. In all cases, data shown are representative of three independent experiments and the error bar represents the standard deviation. Statistical analysis by *t*-test compared with CK (**a**), or the movement of corresponding strains in the absence of *Glycocaulis* 6B-8^T^: *, *p* < 0.05 and **, *p* < 0.01.

## Discussion

Here, we showed that interactions with other species can trigger canonically non-motile bacteria to move on surfaces. Our results clearly demonstrate that this interaction was not derived through a previously reported ‘hitchhiking’ strategy (6). Importantly, apart from initial physical association of the two populations, the surface motility of the sessile species also required its partner pre-colonized the outward region of the starting point at sufficiently high density, thus also possesses different underly mechanism against the interaction between *Pseudomonas fluorescens* Pf0-1 and *Pedobacter* sp. V48 (11). Moreover, we only observed the surface motility under specific environmental conditions, because the presence of biosurfactants was another essential requirement. Therefore, we proposed a previously unknown interaction mode that can promote the migration of canonically non-motile bacteria.

In order to adapt to surface-associated environments, bacteria have evolved a wide array of modes to move along surfaces, most of which require special active appendages, including flagella, pili and fimbriae (1, 3, 5). The genome of strain 45-1b lacks any genes known to encode such structures, suggesting that its migration cannot be attributed to those structure-dependent movement types, such as swarming and twitching. However, one type of surface motility mode that was earlier characterized as appendage-independent is called sliding. Henrichsen *et al* (1972) defined sliding as a passive bacterial translocation created by expansive forces accelerated by surfactants that reduce surface tension (3). The lack of active appendages in strain 45-1b suggests that strain 45-1b exhibited a sliding-like motility in this interaction mode. This hypothesis is also in agreement with the fact that surface motility of 45-1b cells required biosurfactants. Similar phenomena have been reported in *Pseudomonas fluorescens* SBW25 (15) and *Pseudomonas aeruginosa* PAO1 (16), where biosurfactants were essential to aid sliding across a surface. In addition, sliding motility commonly shapes dendritic colony morphology on agar plates (17), a typical feature of our newly described interaction mode.

Interestingly, the putative sliding-like motility of the strain 45-1b is required to be triggered by interaction with the strain 6B-8. We assume that the role of 6B-8 cells in this interaction is related to its contribution to alter the surface properties of the 45-1b cells. Hölscher *et al* (2017) recently expanded the definition of sliding by emphasizing that, in addition to biosurfactants, other cell constituents are possibly involved in reducing surface tension and increasing expansive forces (17). Some reports showed that certain strains of bacteria possess strategies to alter the properties of their own cell surfaces (17–20). For example, *Mycobacterium* synthesize glycopeptidolipids (GPLs), molecules that are part of the outer layer of its capsule, to increase the hydrophobicity of their own cell surface, thus facilitating its sliding on hydrophilic surfaces (19, 20). In SEM images, we found that 6B-8 cells use stalks to attach to the cell surface of 45-1b cells. On the basis of these findings, we hypothesize that such cell-cell contacts change the surface property of 45-1b cells, thus inducing their collective movement. As 6B-8 cells fail to show active motility along surfaces, the migration of 45-1b cells requires sufficiently high numbers of 6B-8 cells pre-colonizing outward region to sustain this beneficial surface state of 45-1b. Furthermore, as this change is based on cell-cell contacts, metabolite exchanges are presumably not sufficient to induce cell motility. Although conceptually reasonable this hypothesis must be further tested in future experiments.

It would be also interesting to look deep into whether the stalk-attachment behavior of 6B-8 cells benefit themselves. Firstly, this behavior should allow 6B-8 cells to stay close with their interaction partners, thus occupying positions exhibiting high metabolite concentration. Secondly, stalks might be used by cells as a nutrient antenna, thus promoting the uptake and transport of the metabolites from the partner cell. A number of relevant clues have been observed in other dimorphic prosthecate bacteria, which have been shown to lengthen their stalks during phosphate starvation, thus accelerating the uptake and transport of phosphate required for growth (21–24). In addition, we found that 6B-8 cells held longer stalks on surface than on liquid environment. As the liquid culture with shaking provided an environment with homogeneous nutrient distribution, this result suggests that when interacting with strain 45-1b on surface, the extracellular metabolites gradients generated by 45-1b cells can induce 6B-8 cells lengthening their stalks, allowing 6B-8 cells attach to and stay close with 45-1b cells.

Dimorphic prosthecate bacteria, including 6B-8, are often found to survive under oligotrophic conditions (25–28). For example, *Caulobacter*, the most studied DPB, fails to grow in rich nutrient media such as LB. In order to adapt to oligotrophic environments, *Caulobacter* has to evolve to exhibit the physiological properties of oligotrophs. Although 6B-8 was also found in extreme and oligotrophic environments, it grows in LB but only utilized a small number of carbon sources (13). Based on these findings, we propose that 6B-8 cells survive in such environments by way of interacting with other bacteria, such as hydrocarbon-degrading bacteria in environments. In order to adapt to an oligotrophic environment, *Caulobacter* has to evolve into an oligotroph. In contrast, 6B-8 has developed the ability to scavenge nutrients from other bacteria. In exchange, 6B-8 assisted other bacteria in moving more freely along surfaces. Movement of these bacteria helped themselves to obtain more nutrients and produce more metabolites, which are in turn made available to 6B-8.

Together, our analysis revealed that a ‘reciprocity’ process is possibly widespread in many habitats, and presumably represents a prevalent ‘win-win’ strategy for bacteria to adapt to perpetually changing environments.

## Methods

### Bacterial strains and culture conditions

The bacterial strains used in this study are described in Table 1. *Dietzia* strains were grown in GPY medium (10 g/L glucose, 10 g/L tryptone, 5 g/L yeast extract) at 30°C. *Caulobacter crescentus* CB15 were grown on PYE medium (2 g/L tryptone, 1 g/L yeast extract, 0.3 g/L MgSO_4_∙7H_2_O, 0.0735 g/L CaCl_2_∙2H_2_O) at 30°C. Other strains were grown in lysogeny broth (LB, 10 g/L tryptone, 5 g/L yeast extract, 10 g/L NaCl, pH=7.0) medium at 30°C (except *E. coli* DH5α, at 37°C). All strains were cultured while shaking at 150 rpm. Both strain 45-1b and 6B-8 were grown in GPY medium and LB medium to their later logarithmic phase respectively, before cells were harvested by centrifugation at 1,500×g, at 4°C for 5 min. Cells were then washed three times with minimal medium (29) (pH=8.0), and re-suspended in minimal medium to prepare cell suspensions for inoculation (OD_600_=5.0). The inocula for other bacterial strains were prepared using the same protocol.

**Table 1.**
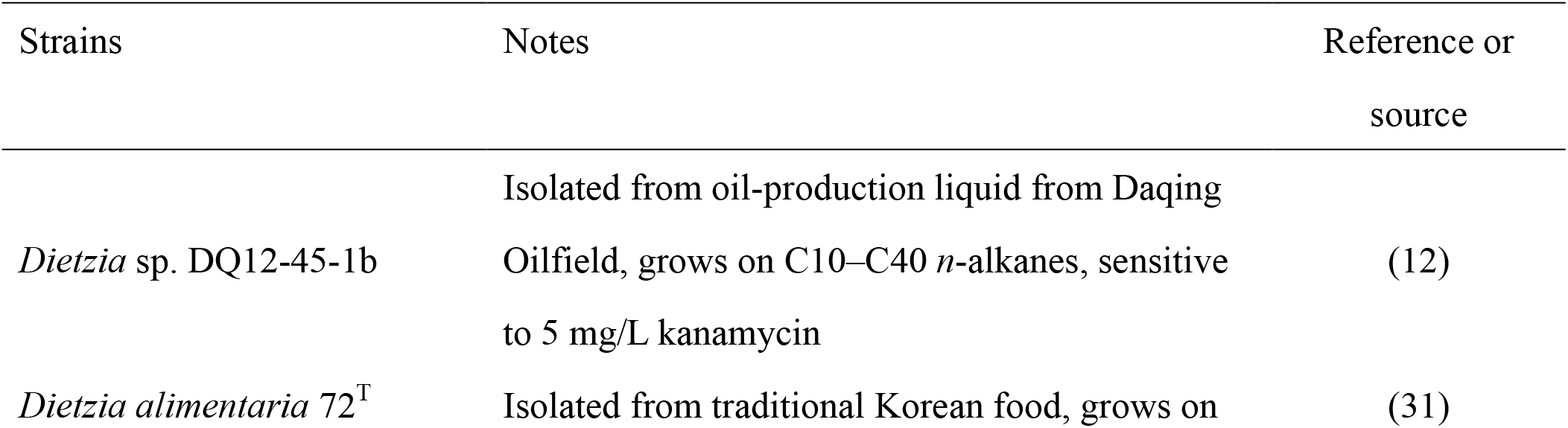
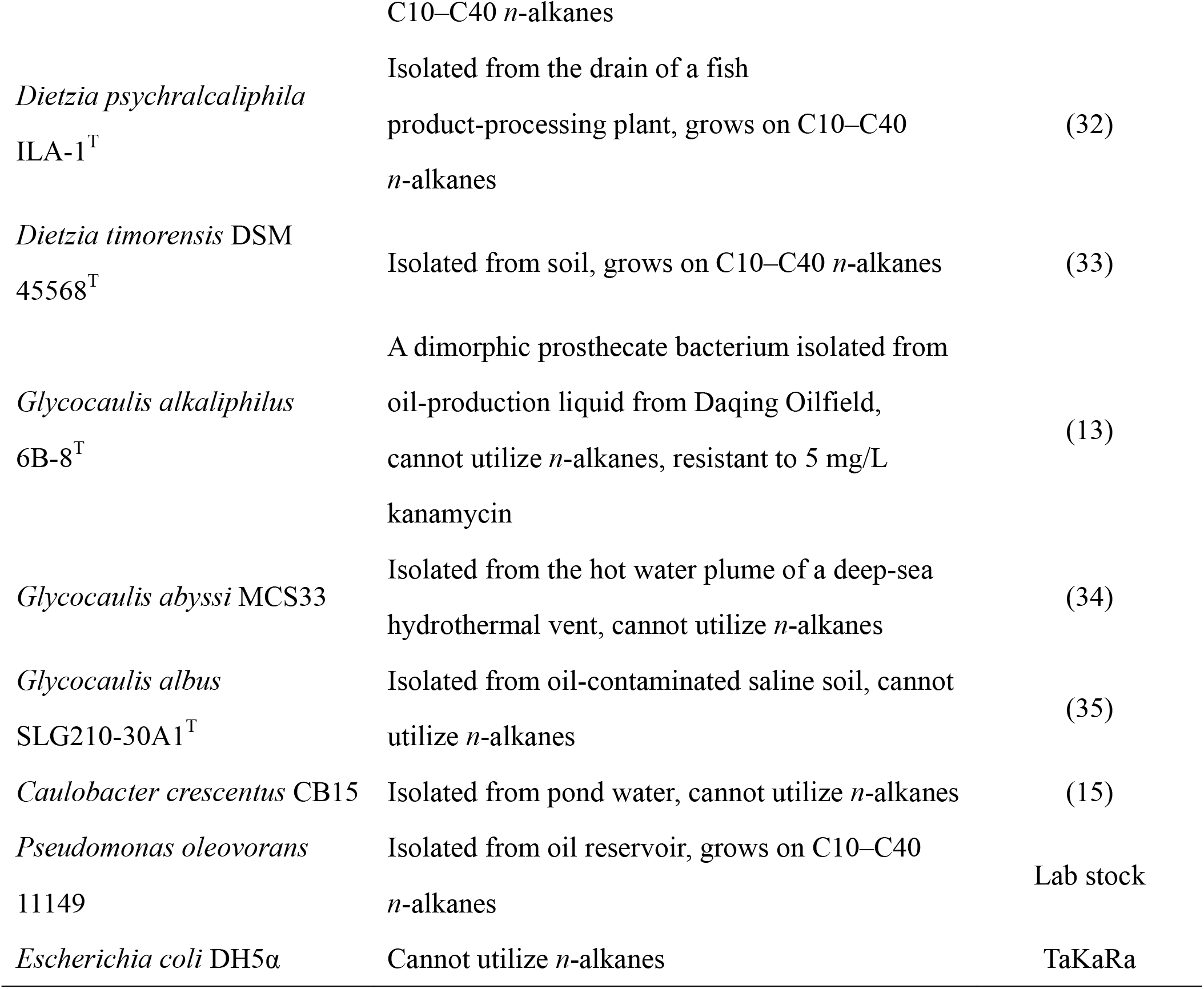
Strains used in the study.

**Table 2.**
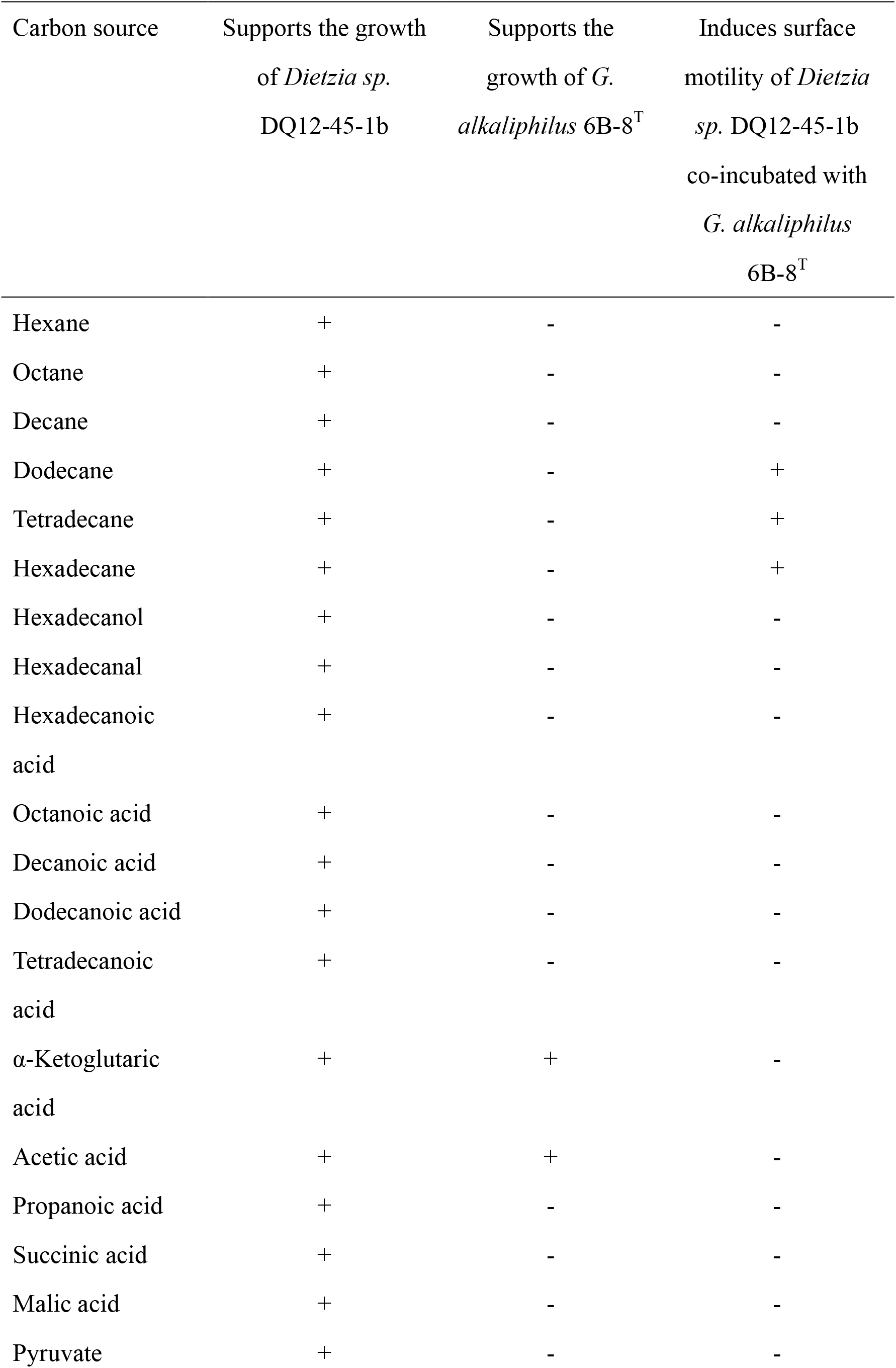
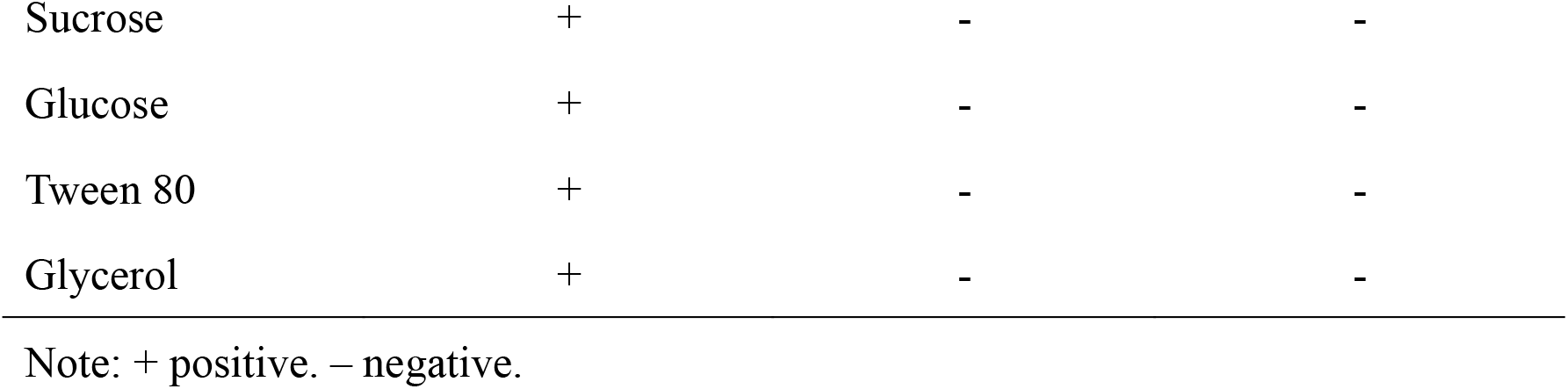
Carbon sources used for the surface motility assay.

| Carbon source | Supports the growth<br>of *Dietzia* sp.<br>DQ12-45-1b | Supports the<br>growth of *G.*<br>*alkaliphilus* 6B-8<sup>T</sup> | Induces surface<br>motility of *Dietzia*<br>sp. DQ12-45-1b<br>co-incubated with<br>*G. alkaliphilus*<br>6B-8<sup>T</sup> |
| --- | --- | --- | --- |
| Hexane | + | - | - |
| Octane | + | - | - |
| Decane | + | - | - |
| Dodecane | + | - | + |
| Tetradecane | + | - | + |
| Hexadecane | + | - | + |
| Hexadecanol | + | - | - |
| Hexadecanal | + | - | - |
| Hexadecanoic<br>acid | + | - | - |
| Octanoic acid | + | - | - |
| Decanoic acid | + | - | - |
| Dodecanoic acid | + | - | - |
| Tetradecanoic<br>acid | + | - | - |
| α-Ketoglutaric<br>acid | + | + | - |
| Acetic acid | + | + | - |
| Propanoic acid | + | - | - |
| Succinic acid | + | - | - |
| Malic acid | + | - | - |
| Pyruvate | + | - | - |
| Sucrose | + | - | - |
| Glucose | + | - | - |
| Tween 80 | + | - | - |
| Glycerol | + | - | - |

Note: + positive. – negative.

### Preparation of soft agar plates

The soft agar plates for surface motility assays were prepared from warm (45°C) minimal medium containing 0.5% agar (Biodee, China, pH=8.0) supplied with different carbon sources. The soluble carbon sources (Table 2) were dissolved in (final concentration of 10 g/L) the agar medium directly. For *n*-alkanes, one piece of sterile filter paper (Grade 2, Whatman, USA; 10 cm in diameter) was fixed inside the lid of a petri dish, followed by addition of 500 μL of the respective *n*-alkane on the filter paper. When the filter paper was saturated with the *n*-alkanes, the lids were sealed with the bottom part of the petri dish that already contained both the agar medium and the inoculated bacteria. To supply *n*-alkanes as carbon sources for bacterial growth, the petri dishes were incubated upside down with the vaporized *n*-alkanes rising to the agar medium. Unless stated otherwise, all plates were incubated in the dark at 30°C for four days.

### Surface motility induced by interactions between the two strains

#### 1) Surface motility assay on agar plate premixed with 6B-8 cells

The soft agar plates pre-mixed with 6B-8 was made by mixing 0.5% agar medium at 45°C with a 6B-8 cell suspension, to obtain agar plates with final 6B-8 concentrations of OD_600_=0.05. Once the agar was solidified, 6B-8 cells were evenly distributed across the agar. Unless stated otherwise, this initial density of 6B-8 was set for all other surface motility assays. To obtain the suspension of dead 6B-8 cells, 6B-8 cell suspensions (OD_600_=5.0) were boiled for 20 minutes and then mixed with agar medium as described above. Subsequently, 1 μL of 45-1b cell suspension was inoculated on the center of plates. 45-1b cells incubated on agar without 6B-8 were used as control. Next, agar plates were incubated with C_16_ as the sole carbon source at 30°C for 4 days.

#### 2) Surface motility assay on agar plate with both cell types initially mixing together

For experiments investigating whether the observed motility derived from a ‘hitchhiking’ strategy, the mixture of 45-1b and 6B-8 suspensions was incubated on the soft agar plates. Briefly, suspension of 45-1b and 6B-8 (OD_600_=5.0) were mixed in equal proportions and inoculated on the center of an agar plate. The agar plates were incubated using C_16_ as the sole carbon source.

#### 3) Surface motility assay on agar plate overlaid with 6B-8 cells

The agar plate overlaid with 6B-8 was prepared by either spreading 100 μL inoculum of 6B-8 cell suspension (OD_600_=1.0) onto one half of the agar plate, or by spreading 20 μL cell suspension on the plate to form a cross pattern, before air-drying for 20 minutes. One μL of 45-1b suspension was placed onto the center of the agar plates. The agar plates were incubated using C_16_ as the sole carbon source.

#### 4) Effect of initial spatial distribution of 6B-8 on the surface motility of 45-1b

One μL of 45-1b suspension (OD_600_=5.0) was inoculated in the center of a soft agar pre-mixed with different initial concentrations (OD_600_=0.001, 0.0015, 0.0025, 0.01, 0.02, 0.1, and 0.2) of 6B-8 cells and using C_16_ as the sole carbon source. A cubic lattice model was used to calculate the average distances between the pre-mixed 6B-8 cells in the agar plate. Assuming that the space containing 6B-8 can be divided into same lattices and the cells evenly distribute in the vertices of a lattice, the side length of the lattices was taken as the average distance between 6B-8 cells. If 1 mL soft agar is divided into a *n×n* cubic lattice, it consists of *n^3^* lattices with a side length of *1/n* cm and totally (*n+1)^3^* lattice vertices, thus with a cell density of *X* cells per mL, *n* can be calculated by *n=X^1/3^*−1, and the average distance can then be obtained: 1/*(X^1/3^*−1) cm. *X* was measured by calibrating the OD_600_ values and colony forming units formed on 1.5% LB agar plates.

#### 5) Surface motility assay using Transwell plates

To measure whether the metabolic exchange between 45-1b and 6B-8 triggers surface motility, Transwell plates (Corning, USA) containing 12 wells were used, each of which contains two chambers and a membrane with pore size of 0.4 μm in diameter. The membrane allows metabolites to be exchanged through the pores while ensuring that the two bacterial strains are kept separate. Fifty mL of warm 0.5% agar medium (at around 45°C) was thoroughly mixed with 6B-8 suspension (500 μL, OD_600_=5.0), and 1 mL of this mixture was pipetted into the outer chamber. Next, 1 mL warm agar medium (at around 45°C) was pipetted into the inner chamber. After the agar medium in both the chambers had solidified, 1 μL of 45-1b cell suspension (OD_600_=5.0) was dropped onto the center of the agar plate in the inner chamber (Fig. 3a). A piece of sterile filter paper (Grade 2, Whatman, USA; approximately 12 cm× 8 cm) was fixed inside the lid of a Transwell plate, saturated with 500 μL C_16_, and then sealed with the Transwell plate. This experiment was performed with both negative and positive controls. For the negative control, 6B-8 was omitted either in the inner or outer chamber of the Transwell plate. In the positive control, the inner chamber agar medium was pre-mixed with 6B-8, while the outer chamber agar medium was not pre-mixed (Fig. 3a). To check the exchange of metabolites, 1 μL fluorescein solution (1 μg/ mL, Sigma, USA) was added on the center of the agar plate in the inner chamber (Fig. 3a), which was inoculated at 30°C for 1 hour and then photographed using a UV transilluminator (Bio-Rad, USA).

#### 6) Surface motility assay on agar plate with separate inoculation on surface

One μL of 6B-8 suspension was inoculated (OD_600_=5.0) on one end of an agar plate slide and strain 45-1b (1 μL, OD_600_=5.0) dropped at two different places, *i.e.*, 1 cm and 2 cm away from 6B-8, which was incubated with C_16_ as the sole carbon source. To verify diffusion of metabolites on the agar, 1 μL fluorescein solution (1 μg/ mL, Sigma, USA) was simultaneously added into the center of the agar plate.

#### 7) Surface motility assay on agar plate overlaid with metabolite-containing supernatant

The metabolite-containing supernatant was prepared using the following procedure. We inoculated both 45-1b and 6B-8 cells into 150 mL minimal medium containing 5 mL/L C_16_ as the sole carbon source, before incubating at 30°C in the dark while shaking for 10 days at 150 rpm. The cultures were centrifuged at 1500 × g, 4°C for 5 min and the supernatant was filtered with a 0.22-μm membrane to remove cells. 200 μL of this supernatant was evenly spread onto an agar plate without 6B-8 and air dried for 20 minutes, 1 μL of 45-1b cell suspension (OD_600_=5.0) was inoculated onto this agar plate and cultured using C_16_ as the sole carbon source. For a negative control, 1 μL of 45-1b cell suspension (OD_600_=5.0) was inoculated onto agar plate overlaid with 200 μL of minimum medium using C_16_ as the sole carbon source.

### Assays on the influence of surfactants surface motility

#### 1) Surface motility assay on agar plate overlaid with surfactant-containing supernatant

Strain 45-1b was inoculated by itself into 150 mL minimal medium containing 5 mL/L C_16_ as the sole carbon source, which was incubated in the dark at 30°C by shaking at 150 rpm for 10 days. The culture was then centrifuged at 1500 × g, 4°C for 5 min, and the supernatant was filtered using a 0.22-μm membrane to remove the cells. 200 μL of this supernatant was evenly spread onto a soft agar pre-mixed with 6B-8 and 10 g/L glucose as the sole carbon source, and air-dried for 20 minutes. 1 μL of 45-1b cell suspension (OD_600_=5.0) was inoculated onto the center of an agar plate.

#### 2) Surface motility assay on agar plate overlaid with rhamnolipid

First, 200 μL of rhamnolipid (Sigma, USA) solution at concentrations of 0.1, 0.5, 1, 5, 50, 100, and 500 mg/L was added onto the soft agar plate premixed with 6B-8 cells and containing glucose (10 g/L) as the sole carbon source. We then added 1 μL of 45-1b cell suspension (OD_600_=5.0) onto the center of the agar plates.

### Assay to screen for potential benefits obtained from interactions between the two strains

#### 1) Growth of 6B-8 on metabolites of 45-1b from degradation of hexadecane

Metabolite-containing supernatants (200 μL, prepared as described above), α-ketoglutarate solution (200 μL, 5 g/L), or a mixture of both (200 μL, volume ratio of 1:1) were spread on an agar plate premixed with a 6B-8 cell suspension, and then incubated at 30°C for four days. Subsequently, a replica plating assay was performed as described below.

#### 2) Surface motility assays with a nutrient gradient

200 μL of surfactant-containing supernatant (prepared as described above) was spread onto a soft agar plate premixed either with or without 6B-8 cells. A strip of sterile paper (4 cm×0.35 cm) saturated with 20 μL of a glucose solution (250 g/L) was placed on one end of the plate. Next, 1 μL of a 45-1b cell suspension was inoculated at a distance of 1 cm, 2 cm, 3 cm, and 3.5 cm away from the paper strip.

##### Screening of different strain combinations for surface motility

We used two combinations of bacterial pairs. First, we inoculated 1 μL of 45-1b cell suspension to the center of soft agar plates pre-mixed with *Glycocaulis abyssi* MCS33, *Glycocaulis albus* SLG210-30A1^T^, *Caulobacter crescentus* CB15, *Pseudomonas oleovorans* 11149, or *E. coli* DH5α (OD_600_=5.0; Table 1) and then incubated the plates in the dark at 30°C for four days, providing C_16_ as the sole carbon source. Second, 1 μL of cell suspensions of *Dietzia alimentaria* 72^T^, *Dietzia psychralcaliphila* ILA-1^T^, or *Dietzia timorensis*, DSM 45568^T^ (Supplementary Table 1) were added in the center of soft agar plates pre-mixed with 6B-8 and incubated with C_16_ as the sole carbon source.

##### Replica plating assay

The colonies grown on the agar plates premixed 6B-8 incubated either with or without 45-1b were blotted using a 0.22-μm sterile membrane (8 cm in diameter) and transferred to a new LB agar plate containing kanamycin (5 mg/L). Replica of the agar plates premixed with 6B-8 and containing metabolites of 45-1b degrading C_16_ was carried out using the same protocol. The blotted LB plates were then incubated for two days at 30°C.

##### Scanning Electron Microscopy Imaging

After 4 days of incubation, 45-1b cells on the soft agar plate overlaid with or without 6B-8 were harvested and washed twice by: using sterile water. In addition, 45-1b and 6B-8 cells were both inoculated at a final cell density of OD_600_=0.05 in 150 mL liquid minimal medium containing C_16_ (final concentration 5 mL/L) as the sole carbon source. Following incubation at 30°C by shaking at 150 rpm for 10 days, the cells were harvested by centrifugation (1500 × g, 5 min, 4°C). The harvested cells were then fixed, dehydrated, and sputter-coated (30), before examination using an SU8010 field emission scanning electron microscope (Hitachi, Japan).

### Statistical and image analysis

Colonies were photographed using a Nikon D5300 camera (Nikon, Japan), and images were analyzed using the Digimizer computer program (MedCalc Software, Version 4.2.6.0). Colony radii of three independent experiments were measured every 24 h. To compare the radii of different sets, we performed unpaired, two-tailed, Student’s t-test using Microsoft Excel for Windows (version 2019, Microsoft Software). To calculate the average movement speed of 45-1b, Colony radii of three independent experiments were measured after 96-h culturing, and speed was calculated by dividing the distance traveled by the time required.

The length of the stalks of 6B-8 attached to 45-1b cells were measured using a Digimizer computer program (MedCalc Software, Version 4.2.6.0) based on SEM images. To compare the stalk length of different culture conditions (liquid or agar surface), we performed unpaired, two-tailed, Student’s t-test, using Microsoft Excel for Windows (version 2019, Microsoft Software).

## Acknowledgments

This work was supported by National Key R&D Program of China (2018YFA0902100 and 2018YFA0902103), and National Natural Science Foundation of China (31770118 to YN, 31770120 and 31761133006 to XLW).

